# Integrin-αVβ3 is a fundamental factor in medulloblastoma tumorigenicity and radioresistance: A new game for an old player

**DOI:** 10.1101/2023.03.08.531652

**Authors:** William Echavidre, Jérôme Durivault, Célia Gotorbe, Thays Blanchard, Marina Pagnuzzi-Boncompagni, Valérie Vial, Florian Raes, Alexis Broisat, Rémy Villeneuve, Régis Amblard, Nicolas Garnier, Cécile Ortholan, Marc Faraggi, Benjamin Serrano, Vincent Picco, Christopher Montemagno

## Abstract

Medulloblastoma (MB) is the most frequent solid tumor in children, localized in the brain’s posterior fossa. Its standard of care comprises maximal resection surgery followed by craniospinal irradiation and chemotherapy. Despite a long-term survival rate of 70%, wide disparities among patients have been observed. Relevant targets for naive and recurrent MB are urgently needed. Primary and recurrent MBs are characterized by aggressive invasion into surrounding brain tissue, active angiogenesis, and radioresistance. Integrin-αvβ3 was a major driver of these features in glioblastoma. Nevertheless, such observations have not yet been reported in MB. Integrin-αvβ3 was found to be expressed in a subset of MB patients. We investigated the role of integrin-αvβ3 using MB-derived cell lines with β3-subunit depletion or overexpression both in vitro and in vivo. Radioresistant MB cell lines were generated and showed increased integrin-αvβ3 expression, which correlated with increased susceptibility to pharmacological integrin-αvβ3 inhibition with cilengitide, a competitive ligand mimetic. Finally, we conducted single-photon emission computed tomography (SPECT)/magnetic resonance imaging (MRI) studies on orthotopic models using a radiolabeled integrin-αvβ3 ligand (^99m^Tc-RAFT-RGD). This approach offers the prospect of a novel predictive imaging modality in MB. Altogether, our data pave the way for SPECT/MRI-based selection of a subpopulation of MB patients eligible for integrin-αvβ3-directed therapies.

**SIGNIFICANCE:** This study demonstrates integrin-αvβ3’s fundamental role in MB tumorigenicity and radioresistance and the effect of its expression on cilengitide functional activity.

## INTRODUCTION

Brain tumors are the second most common childhood tumors after hematologic cancers (1). Medulloblastoma (MB) is the most common brain tumor in the pediatric population, accounting for approximately 20% of all childhood brain tumors (2). MBs arise in the posterior fossa, mainly in the cerebellum, most often during early life (3). They represent about 60% of all embryonal tumors, making them the most common cause of death in pediatric oncology (4,5). The current standard of care for MBs includes a maximal tumor resection followed by craniospinal irradiation and chemotherapy (6). Recent advances in the molecular understanding of MB pathophysiology have defined four prognosis molecular subgroups: wingless (WNT), sonic hedgehog (SHH), group 3, and group 4. Nevertheless, the clinical translation of these observations is just starting to emerge. Therefore, its treatment modalities remain based on classifying patients into standard- or high-risk groups according to the presence of metastases, age, and extent of postoperative residual disease.

Standard-risk patients (70% of MBs) are children aged >3 years with no evidence of disseminated disease and postoperative residual tumor of <1.5 cm^2^. In this case, the five-year post-diagnosis overall survival (OS) is 85%. If at least one of these criteria is unmet, the patient is considered high-risk, and the 5-year OS drops to 60% (7). Despite the recent identification of four core molecular subgroups (SHH, WNT, group 3, and group 4), clinical stratification of the patients still mainly relies on the clinical features described above. In addition, the tumors of about 30% of the patients originally included in the standard-risk group eventually relapse. Therefore, accurate identification of high-risk MBs and management of recurrent MBs remain major challenges since prognostic markers remain lacking, and treatment options are limited in case of recurrence. These issues highlight the need to refine MB diagnostic criteria with new prognostic and predictive markers and therapeutic options designed explicitly for recurring disease.

In the past decades, understanding the molecular pathogenesis of malignant brain tumors such as gliomas has led to the identification of novel regulators of glioblastoma (GBM) cell growth, invasion, and angiogenesis (8). Integrins were identified as major effectors of these processes and proposed as attractive therapeutic targets for GBM. Integrins are a large family comprising non-covalent heterodimeric complexes involving 18 α and eight β subunits, possibly forming at least 24 different receptor types (9). Integrins are membrane-bound receptors specific to interactions between cells and extracellular matrix (ECM) components such as fibronectin, laminin, and collagen. Upon binding to their ligands, integrins activate downstream signaling pathways that regulate many cellular effects under physiological and pathological conditions. While integrin-encoding genes are rarely mutated in cancer, dysregulation of their expression level or integrin signaling is common, especially in brain tumors (10).

Among these integrin complexes, integrin-αvβ3 was the first to be found aberrantly expressed in high-grade brain tumors. Integrin-αvβ3 belongs to the integrin subtypes that recognize the tripeptide Arg-Gly-Asp (RGD) sequence found in many ECM proteins, including fibronectin and vitronectin. Many studies have highlighted the central role of avβ3 in sustaining the proliferation and invasive properties of brain tumors through the activation of focal adhesion kinase (FAK), leading to the activation of downstream pathways, including the phosphoinositide 3 kinase/protein kinase B (Akt) and extracellular signal-regulated kinase (ERK)/mitogen-activated protein kinase pathways (8).

Given their involvement in tumor progression, integrins are considered attractive therapeutic targets for solid tumors. Therefore, several strategies have been explored to target integrin-αvβ3, including small molecules. They include a cyclic pentapeptide blocking the RGD binding site (cilengitide; cyclo-Arg-Gly-Asp-DPhe-NMe-Val) that acts as a selective αvβ3 inhibitor. Cilengitide was found to impair GBM tumorigenesis by hampering angiogenesis, cell proliferation, and invasion. This agent was subsequently evaluated in clinical studies (11,12). Cilengitide had moderate anti-tumor efficacy in GBM patients, with anti-tumor effects restricted to high-avβ3-expressing tumors, highlighting the potential need to stratify patients based on avβ3 expression in their tumors (13). In this context, the advent of nuclear medicine is currently one of the greatest components of theranostics concepts. As a molecular probe, radiolabeled RGD was developed for αvβ3-based imaging and therapy of solid tumors, including GBM (10). Nevertheless, such studies remain to be conducted for MB.

This study aimed to evaluate the role and the relevance of integrin-αvβ3 in MB for the first time. First, the integrin-avβ3 expression was measured in patient-derived samples and naive and radioresistant MB cell lines. Then, integrin-αvβ3’s role was explored using genetic depletion of the β3 subunit or cilengitide in vitro and in vivo in orthotopic MB mouse models. Finally, molecular imaging with ^99m^Tc-radiolabeled-RGD peptides was used to noninvasively measure integrin-αvβ3 expression in MBs.

## MATERIALS AND METHODS

### Cell lines and culture conditions

DAOY (American Type Culture Collection [ATCC]: HTB-186; SHH group) and CHLA-01-Med (ATCC: CRL-3021) were obtained from the ATCC. HD-MB03 (Deutsche Sammlung von Mikroorganismen und Zellkulturen [DMSZ]: ACC 740; group 3) was obtained from DMSZ. D458 (Cellosaurus: CVCL_1161; group 3) was from Dr. Celio Pouponnot (Institut Curie, Paris, France). ONS-76 (SHH group) was obtained from Dr. Gilles Pagès (Institute for Research on Cancer and Aging-Nice, Nice, France). DAOY, D458, and ONS-76 cells were cultured in Dulbecco’s modified eagle medium (DMEM) supplemented with 1 mM sodium pyruvate, 2 mM Glutamax, and 7.5% fetal bovine serum (FBS). CHLA-01-Med cells were cultured in DMEM/F12 supplemented with 2% B-27, 20 ng/mL epidermal growth factor (EGF), and 20 ng/mL basiFGF. HD-MB03 cells were cultured in Roswell Park Memorial Institute medium supplemented with 7.5% FBS. Cells were grown at 37°C with 5% carbon dioxide (CO_2_).

### Genetic disruption of β3-integrin using CRISPR-Cas9, β3-integrin expression rescue, and lentiviral transductions

DAOY wildtype cells were transfected using jetPEI (Polyplus, 101000053) according to the manufacturer’s instructions with pSpCas9(BB)-2A-GFP (PX458) plasmid (a gift from Feng Zhang; Addgene plasmid #48138; http://n2t.net/addgene:48138; research resource identifier: Addgene_48138) containing clustered regularly interspaced short palindromic repeats (CRISPR)-CRISPR-associated protein 9 (Cas9) targeting regions for the first (guide RNA [gRNA]: 5’-GAGGCGGACGAGATGCGAGCG-3’) and tenth (gRNA 5’-AGACGGGCTGACCCTCCCGG-3’) exons of the β3-integrin gene (*ITGB3*). The *ITGB3* sequence (pcDNA3.1-beta-3 was a gift from Timothy Springer; Addgene plasmid # 27289) was subcloned into pLenti6.3/TO/V5-Blasti (A11144, Thermo Fischer Scientific) to create β3-integrin expression plasmids. Lentiviruses were produced by triple transfection of HEK-293T cells with the lentiviral transfer vector pLenti6.3/TO/V5-Blasti and the packaging plasmids psPAX2 (Addgene plasmid #12260) and pMD2.G (Addgene plasmid #12259), a kind gift from Didier Trono, at a ratio of 0.3:0.3:0.1. Transfection was performed using jetPEI (101000053, Polyplus). The viral supernatant was collected 48 h after transfection, filtered through a 0.45 μm filter, and added to the target cells.

For in vivo tumorigenesis assays, DAOY- and HD-MB03-derived cells were transduced with lentiviral particles containing RFP-Luc (Lenti-One RFP-Luc, GEG Tech) at a multiplicity of infection of 0.3 in the presence of hexadimethrine bromide (4 μg/mL).

### RT-qPCR

Real-time reverse transcription-quantitative polymerase chain reaction (RT-qPCR) analyses were performed using human cerebellar mRNA (Biochain) and MB cell lines. Messenger RNAs (mRNAs) were prepared using a Nucleospin RNA Kit (Macherey-Nagel), and complementary DNA (cDNA) synthesis was performed using a Maxima First Strand cDNA Synthesis Kit for RT-qPCR with dsDNase (Thermo Fischer Scientific). Quantitative PCR analyses were performed on an Applied Biosystems StepOnePlu System using TB Green Premix Ex Ta (Tli RNase H Plus; Takara Bio) reagents. The primers used are listed in Supplementary Table S1. Relative expression levels were determined using the ΔCt method and normalized to the reference gene 36B4. Results are expressed relative to the normal cerebellum.

### Immunoblotting

Cells were lysed in 1.5× Laemmli buffer, and protein concentrations were determined using the Pierce BCA Protein Assay (Thermo Fisher Scientific). Protein extracts (40 μg) were separated via electrophoresis on 10% sodium dodecyl-sulfate-polyacrylamide gels and transferred to polyvinylidene difluoride membranes (Millipore). Membranes were blocked in 2% milk-phosphate-buffered saline (PBS) and incubated with the following antihuman antibodies: rabbit β3-integrin (1:1000; 13166, Cell Signaling Technology [CST]), rabbit p-FAK (1:1000; 8566, CST), rabbit FAK (1:1000; 71433, CST), mouse p-Akt (1:1000; 4051, CST), rabbit Akt (1:1000; 9272, CST), rabbit p-ERK1/2 (1:1000; 4370, CST), rabbit ERK1/2 (1:1000; 4695, CST), and rabbit poly (ADP-ribose) polymerase (PARP; 1:1000, 9542; CST). Actin was used as the protein loading control (1:5000; MA5-15739, Thermo Fisher Scientific). Immunoreactive bands were detected with horseradish peroxidase-coupled anti-mouse or anti-rabbit antibodies (CST) using the ECL System (Merck Millipore; WBKLS0500). Immunoblot analysis was performed using the LI-COR Odyssey Imaging System.

### Measurement of integrin-αVβ3 expression by FACS

DAOY- and HD-MB03-derived cells were seeded into six-well dishes for 24 h. Next, cells were rinsed with PBS, detached with accutase (00-4555-56, Thermo Fisher Scientific), and collected. Then, the cells (250,000) were incubated with a mouse anti-αvβ3 antibodies (1 μg/mL; ab190147, Abcam) for 1 h in PBS/1% bovine serum albumin (BSA) on ice. Next, cells were then centrifuged and washed twice with PBS. Then, cells were incubated with a goat anti-mouse IgG secondary antibody coupled to AlexaFluor-488 (1 μg/mL; A-11029, Invitrogen). All experiments were performed in triplicate. Ten thousand events were analyzed per sample using a BD Fluorescence-Activated Cell Sorting (FACS) Melody cytometer (BD Biosciences). Data were analyzed using the FlowJo software.

### Cell adhesion assays

Cell adhesion to ECM proteins was performed using a CytoSelect 48-well Cell Adhesion Assay Kit (Cell Biolabs) according to the manufacturer’s protocol. Briefly, 100,000 DAOY-derived and 250,000 HD-MB03-derived cells were collected in 200 μl of serum-free medium and added to the pre-warmed ECM adhesion plate. The plates were incubated at 37°C for 90 min in a CO2 incubator. The wells were washed three times with PBS and stained with 200 μl of cell staining solution for 10 min at room temperature (RT). After washing and drying, the stained cells were extracted with extraction solution and incubated in an orbital shaker for 10 min. Then, 150 μl of each sample was transferred to a 96-well plate and quantified by measuring absorbance at 560 nm.

To determine cilengitide’s half-maximal inhibitory concentration (IC50), 1 μg of fibronectin was plated onto 96-well plates for 60 min at 37°C in a CO2 incubator. Serum-starved cells were added to the fibronectin-coated wells (10,000 cells for DAOY and 30,000 for HD-MB03) in the presence of serial cilengitide dilutions (0–200 μM), and the plates were incubated for 90 min at 37°C with 5% CO_2_. Then, cells were washed three times with PBS and stained for 20 min with a 1% Crystal Violet solution at RT. After washing with PBS, adherent cells were solubilized with dimethylsulfoxide (DMSO) and quantified by measuring absorbance at 590 nm. The IC50 was calculated using the nonlinear regression method in the Prism 8 software (Graphpad Software Inc.).

### Proliferation assay

The different cell lines (25,000 and 50,000 DAOY- and HD-MB03-derived cells, respectively) were seeded onto six-well plates in triplicate per cell line and per condition. Cell proliferation was measured by daily trypsinization and counting (Coulter Z1; Beckman) for four days (96 h). The cell proliferation index was calculated as the percentage of day 0 by standardizing each measurement to the cell number obtained 24 h after seeding (day 0) or after initiation of cilengitide treatment (0.5 or 2.5 μM).

### MTT assay

The different cell lines (3,000 and 10,000 for DAOY and HD-MB03 cells, respectively) were seeded in 96-well plates (Corning Inc.) in 100 μl of medium per well. Cilengitide concentrations from 0 to 500 μM were tested. Its effect was measured using the 3-(4,5-dimethylthiazol-2yl)-diphenyltetrazolium bromide (MTT) colorimetric assay (Sigma-Aldrich) according to the manufacturer’s instructions. When applicable, the half-maximal effective concentration (EC50) was calculated using the nonlinear regression method in the Prism 5 software (Graphpad Software Inc.).

### Migration and invasion assays

In vitro migration and invasion assays were performed in Boyden chambers (Corning) with porous membranes coated or not with 50 μg/mL Matrigel for 2 h at 37°C to assess invasion or migration, respectively. Next, DAOY (25,000 cells) and HD-MB03 (75,000 cells) cells were placed in serum-free DMEM in the upper compartment (with or without cilengitide at 0.5 or 2.5 μM) and DMEM in the lower compartment of the chambers. After 12 h and 24 h incubation for DAOY and HD-MB03, respectively, cells were stained with Giemsa for 30 min. Invasive cells were counted under the microscope. Results are expressed as % of the control.

### Cell death

Cells were seeded in 12-well plates (50,000–150,000 cells per well, triplicate per condition) at 37°C/5% CO2 in their respective media. Cells were treated with cilengitide (20 μM) for 48 h. Next, both floating and adherent cells were collected and centrifuged. Then, cell pellets were resuspended in FACS buffer (PBS, 0.2% BSA, and 2 mmol/L ethylenediaminetetraacetic acid) and stained with 2 μg/mL propidium iodide (PI; Invitrogen). PI was added just before the analysis. Experiments were performed at least three times, 10,000 events were analyzed per sample using a BD FACSMelody cytometer, and data were analyzed using the FlowJo software.

### Generation of radioresistant cells

Cells were grown to approximately 50% confluence in 100 mm Petri dishes. Cells were irradiated with 8 Gy of X-rays using a 6 MeV Novalis TrueBeam linear accelerator (Novalis TrueBeam STX, 3 Gy/min) and then returned to the incubator. When they reached about 90% confluence, the cells were trypsinized and subcultured into new dishes. The cells were irradiated again when they reached approximately 50% confluence (second fraction). Fractionated irradiations were continued until the total dose reached 64 Gy. Cells were considered radioresistant if they survived these cycles. Control cells (naïve) were subjected to identical trypsinization, replating, and culture conditions but were not irradiated. For all assays on irradiated cells, there was an interval of ≥2 weeks between the last irradiation and the experiment.

### Immunofluorescence

Tumor sections (5 μm cryostat sections) were fixed in 4% paraformaldehyde for 10 min at RT and blocked in 1% horse serum in Tris-buffered saline (TBS) for 1 h. Next, the sections were incubated with rat monoclonal antimouse platelet and endothelial cell adhesion molecule 1 (PECAM1/CD31; clone MEC 13.3, 1:1000; BD Pharmingen) and monoclonal anti-mouse α-smooth muscle actin (αSMA; A2547, 1:1000; Sigma) or anti-Ki67 (ab16667, 1:500; Abcam) antibodies diluted in TBS containing 1% BSA at 1:100 overnight at RT, and then washed with TBS containing 0.025% Triton. Then, the preparations were incubated with anti-rabbit Alexa488- (#4412, CST) and anti-mouse Alexa-555-coupled secondary antibodies. Next, preparations were washed with TBS containing 0.025% Triton, and the nuclei were counterstained with Hoechst33342 (Thermo Fisher Scientific). Fluorescence images of the cells were taken with a DMI400 (Leica Microsystems) inverted microscope equipped with a 10× objective (Leica Microsystems) and a Zyla 5.5 camera (Andor Technologies). Preparations were mounted and imaged using a Leica microscope (DMI4000B, Leica) and counted at 10× (CD31/αSMA) or 40× (ki67) magnification with Fiji software (14)

### IHC on MB patient samples

Integrin-αvβ3 expression in brain MB sections was assessed by immunohistochemistry (IHC; Tissue Microarray, Biomax). Briefly, after de-paraffinization, sections were saturated with TBS containing 1% BSA and 1% horse serum for 1 h at RT. Next, tumor sections were incubated with a mouse anti-human anti-integrin-αvβ3 antibody (1:100; ab7166, Abcam) overnight at 4°C. Then, sections were washed with TBS containing 0.25% Triton and incubated with a secondary biotinylated antibody (Vectastain ABC-HRP Kit, peroxidase, PK-4002, vector), followed by 3,3’-diaminobenzidine stain (Vector).

### Intracranial orthotopic tumor xenograft models

DAOY-Luc spheroids (eight per animal) or HD-MB03-Luc cell suspensions (5,000 cells per animal) were stereotaxically implanted into the brains of nine-week-old Rj:NMRI-Foxn1 nude (nu/nu) female mice (Janvier Labs). Briefly, DAOY spheroids were generated with 2500 cells grown for 48 h in ultralow-adhesion spheroid 96-well plates (Corning). DAOY-Luc spheroids and HDMB-03-Luc cells were implanted into the left cerebellar hemisphere (2 mm posterior, 1.5 mm left of the lambda point, and 2.5 mm deep) using a Hamilton syringe fitted with a needle (Hamilton) and following a previously described procedure (15). Cilengitide was resuspended in 200 μL of an aqueous solution of 5% DMSO in PBS. Mice were administered 300 μg cilengitide three times per week via intraperitoneal (IP) injection. IP with vehicle solution served as a control. Survival of mice was evaluated by daily monitoring using neuropathological symptoms, including gait abnormalities and weight loss >10% as endpoints. At least six mice per group were selected to achieve sufficient statistical power.

### Bioluminescence imaging

Mice were imaged two days after cell implantation, and tumor growth was evaluated by bioluminescent imaging (BLI; IVIS, PerkinElmer). Mice were monitored for up to 170 days (DAOY-derived cells) or 50 days (HD-MB03-derived cells) after IP injection of 3.3 mg of D-Luciferin dissolved in 100 μL of PBS. Ten min after injection, mice were anesthetized with isoflurane and imaged with an IVIS Spectrum (field of view: C; binning: medium, f-stop: 1; exposure time: 1 min). Bioluminescence signals (total flux [photons/sec]) were quantified using Living Image 2.0 (Caliper Life Sciences).

### SPECT/MRI imaging and autoradiography

The in vivo imaging studies used 12 nine-week-old Rj:NMRI-Foxn1 nude (nu/nu) female mice (Janvier Labs). Acquisitions were made using dedicated magnetic resonance imaging (MRI) and single-photon emission computed tomography (SPECT)/computed tomography (CT) MRI systems on six mice bearing DAOY tumors and six mice bearing HD-MB03 tumors seven and four weeks after tumor implantation, respectively, (Nanoscan PET/MRI3T and Nanoscan SPECT/CT; Mediso). Brain tumors were visualized using T1-weighted coronal images acquired after shimming and 2 min after intravenous administration of 100 μL of DOTAREM (0.5 mmol/mL gadoteric acid) using a fast spin-echo MRI sequence (repetition/echo time = 526/27.8 ms, 12 average, 32 slices, and voxel size = 0.34×0.34×1 mm^3^). Once tumors were observable on MRI acquisitions, SPECT/CT acquisitions were performed 1 h after intravenous injection of 63.0 ± 12.6 MBq of ^99m^Tc-RAFT-RGD. SPECT quantification was performed using volumes of interest located within the tumor and contralateral healthy cerebellum, and the tumor-to-contralateral ratio was determined (VivoQuant, Invicro). Then, the animals were euthanized using CO2 inhalation, and the brain was harvested. Following ex vivo gamma-well counting, left and right cerebellum samples were frozen, and 20 μm thick slices were obtained (NX50V, MM France). After overnight exposure to a phosphorscreen (BAS-IP SR 2025 E, Cytiva), autoradiographic images were acquired (Typhoon IP, Cytiva) and quantified using dedicated software (ImageQuantTL). Regions of interest were drawn around tumor lesions and healthy contralateral cerebellum on a minimum of three slices. Results were background corrected and expressed as average tumor-to-contralateral ratios.

### Study approval

All animal experiments were conducted in strict accordance with the recommendations of the Guide for the Care and Use of Laboratory Animals. All animal studies were approved in advance by the local animal care committee (Veterinary Service and Direction of Sanitary and Social Action of Monaco; APAFIS # 19480-2019022616164184v4).

### Statistical analysis

Data are presented as mean ± standard error of the mean (SEM). Data were compared between groups using non-parametric Mann–Whitney (two groups) or two-way analysis of variance (ANOVA) corrected for multiple comparisons using Sidak’s test (>2 groups). Tumor growth was compared between groups using two-way ANOVA corrected for multiple comparisons using Sidak’s test. Survival was compared between groups using the log-rank test (Mantel–Cox). A *p* < 0.05 was statistically significant.

## RESULTS

### Integrin-αvβ3 expression in MBs

Integrin-αvβ3 expression in MB-derived patient samples was first assessed by IHC (Figure 1A). Among the 20 patients analyzed, 20% (4/20) expressed significant integrin-αvβ3 levels. Since several integrins are upregulated in brain tumors, their expression was examined in several MB cell lines (ONS-76, DAOY, D458, CHLA-01Med, and HD-MB03). Most integrin genes were overexpressed in DAOY and ONS-76 (SHH group) cell lines but showed low expression in HD-MB03, D458, and CHLA-01 (groups 3 and 4) cell lines (Figure 1B). Among them, high *ITGB3* expression was detected in DAOY and ONS-76. This observation was confirmed at the protein level with higher expression in DAOY cells (Figure 1C). To investigate the role of αvβ3 in MB tumorigenesis, *ITGB3* was knocked out (KO) in DAOY cells using CRISPR/Cas9 technology, and β3-was overexpressed in DAOY-KO and HD-MB03 cells (Figures 1D-E and S2).

**Figure 1.**
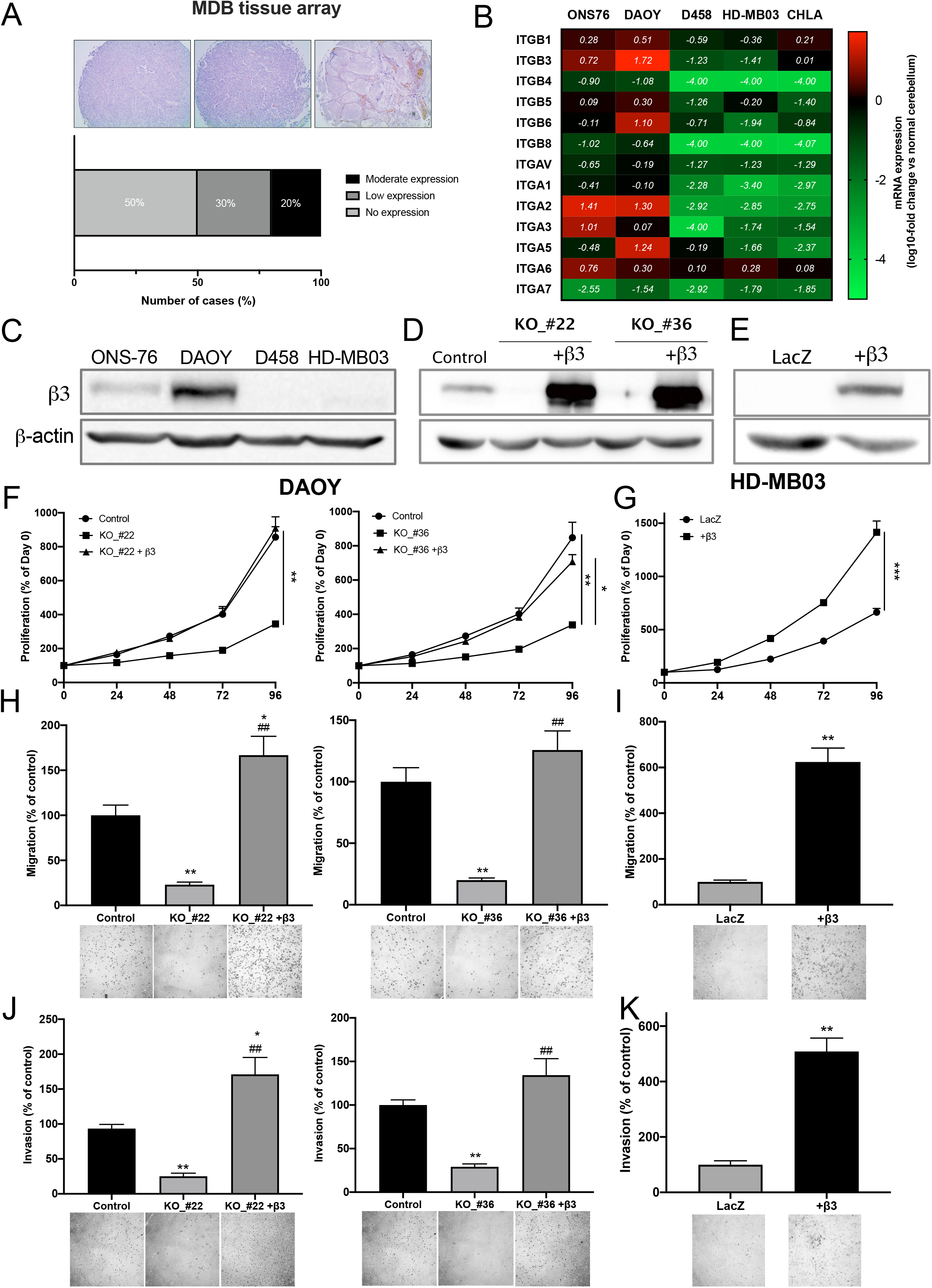
Integrin-αvβ3 is differentially expressed in MB cell lines and promotes their proliferation, migration, and invasion. (**A**) IHC analysis of integrin-αvβ3 expression in MB tumor samples (*n* = 20). Staining intensity was classified as no, low, or moderate expression. (**B**) A heatmap showing integrin gene expression in MB cell lines quantified by RT-qPCR. Results are expressed in comparison to normal cerebellum (log_10_-fold change). The mean values of three independent experiments are shown. (**C**) A western blot of β3-integrin protein expression in MB cell lines. (**D**-**E**) Representative western blots of β3-integrin expression in DAOY (control vs. KO and β3-integrin-overexpressing cells) and HD-MB03 (LacZ vs. β3-integrin-overexpressing cells) cell lines. Three independent experiments were performed. (**F-G**) The proliferation of DAOY_Control, KO_#22, KO_#22-rescue (F, left panel), KO_#36, KO_#36-rescue (F, right panel), and HD-MB03-LacZ and overexpressing-β3-integrin (G). Proliferation rates represent the percentage change vs. day 0. Three independent experiments were performed, and data are presented as mean ± SEM. Key: **, *p* < 0.01; ***, *p* < 0.001. (**H**-**I**) The migration of DAOY_Control, KO_#22, KO_#22-rescue (H, left panel), KO_#36, KO_#36-rescue (H, right panel), and HD-MB03-LacZ and overexpressing-β3-integrin (I). Serum-starved cells were allowed to migrate for 12 h (DAOY) or 24 h (HD-MB03). Migrations were performed using Boyden chamber assays, and the results are presented as the percentage of control cells. Three independent experiments were performed, and data are expressed as mean ± SEM. Key: **, *p* < 0.01 vs. control; ##, *p* < 0.01 vs. KO_cells. (**J**-**K**) The invasion of DAOY_Control, KO_#22, KO_#22-rescue (J, left panel), KO_#36, KO_#36-rescue (J, right panel), and HD-MB03-LacZ and overexpressing-β3-integrin (K). Serum-starved cells were allowed to invade for 12 h (DAOY) or 24 h (HD-MB03). Invasions were performed in a Boyden chamber coated with Matrigel, and results are presented as the percentage of control cells. Three independent experiments were performed, and data are presented as mean ± SEM. Key: *, *p* < 0.05; **, *p* < 0.01 vs. control; ^##^, *p* < 0.01 vs. KO_cells.

### Integrin-αvβ3 promotes MB cell proliferation, migration, and invasion

Because a major function of integrin-αvβ3 is cell adhesion to the ECM, an adhesion assay was first performed on various ECM proteins (Figure S3). Genetic ablation of the β3-subunit in both KO-clones #22 and #36 resulted in significantly decreased adhesion of DAOY cells to fibronectin and fibrinogen. Overexpression of the β3-subunit in DAOY-KO clones or HD-MB03 cells restored cell adhesion to these ECM proteins (Figure S3). The proliferation of DAOY-KO cells was significantly slower than control and β3-overexpressing cells (Figure 1F). The same results were observed with HD-MB03 cells (Figure 1G). Analysis of downstream-integrin-αvβ3 signaling revealed decreased pFAK, pAkt, and pERK1/2 levels in DAOY-KO and HD-MB03_WT cells (Figure S4). As a key player in cell adhesion, the role of integrin-αvβ3 in migration and invasion processes was further investigated. Genetic ablation of the β3-subunit decreased cell migration of both KO clones by 80% (Figure 1H), while its overexpression in HD-MB03 cells increased cell migration by 600% (Figure 1I). Similar observations were made in invasion assays (Figure 1J-K). Restitution of β3-integrin in DAOY-KO cells completely abolished the anti-migratory and anti-invasive effects of β3-depletion. Therefore, integrin-αvβ3 promotes MB cell proliferation, migration, and invasion.

### Integrin-αvβ3 promotes tumorigenesis of orthotopic MB xenografts

The fundamental role of integrin-αvβ3 in tumorigenesis features prompted us to investigate its role in vivo in orthotopic MB models. β3 depletion in DAOY cells impaired orthotopic tumor growth (Figure 2A-C), increasing the median survival of mice bearing DAOY-KO_#22 and KO_#36 tumors by 95% and 54%, respectively, compared to control (*p* < 0.001 vs. DAOY_control; Figure 2E). In HD-MB03 xenografts, β3 expression increased tumor growth rate and decreased mouse survival by 20% compared to control tumors (Figures 2B, 2D, and 2F). Consistent with these observations, ex vivo analyses show an increase in Ki67-positive nuclei in β3-expressing DAOY or HD-MB03 tumors (Figure 2G-H). Altogether, these results highlight the pro-tumorigenic role of integrin-αvβ3 in MB.

**Figure 2.**
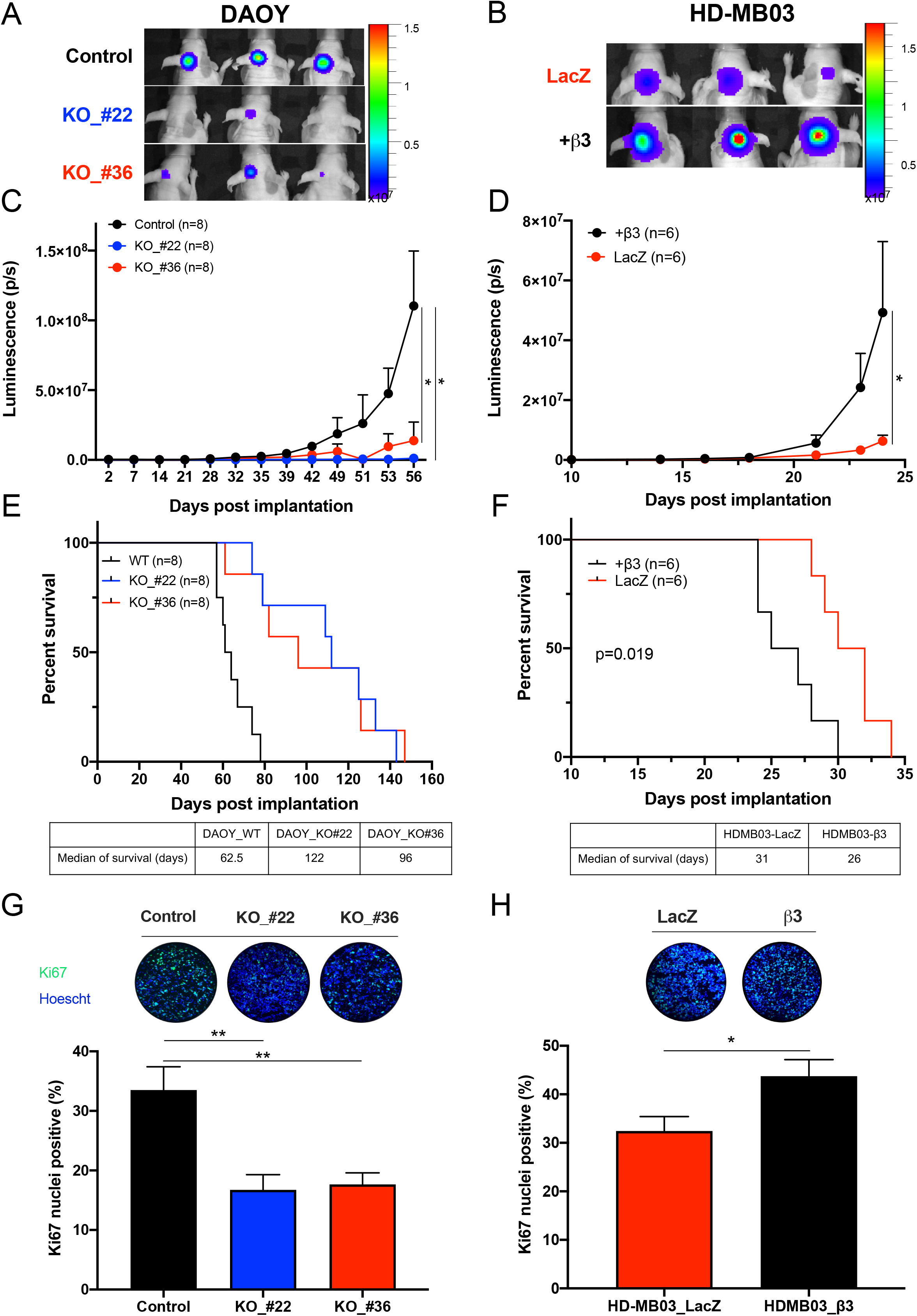
Integrin-αvβ3 promotes MB intracranial tumor growth rate and increases survival. (**A-B**) Representative images of BLI signal intensity 56 days (DAOY_control vs. KO_#22 and KO_#36; A) and 23 days (HD-MB03-LacZ vs. β3+; B) after implantation. Three mice are shown per condition. (**C-D**) Tumor growth of DAOY_control compared to KO_#22 and KO_#36 (**C**) and HD-MB03-LacZ compared to β3+ (**D**) measured by luciferase activity. Photon flux was quantified and analyzed using the IVIS imaging system. Key: *, *p* < 0.05 vs. control. (**E-F**) Survival curves of mice orthotopically implanted with DAOY_control vs. KO_#22 and KO_#36 (**E**) and HD-MB03-LacZ vs. β3+ (**F**) in the cerebellum. Day 0 corresponds to the day of tumor implantation. The median of survival is shown at the bottom of the graphs. The *p*-value is indicated in the graph (Log-rank test). (**G-H**) Ki67 labeling of DAOY_control vs. KO_#22 and KO_#36 (**G**) and HD-MB03-LacZ vs. β3+ (**H**). Representative images of Ki67 immunostaining (green) and Hoechst33342 nuclear DNA counterstaining (blue) are shown at the top of the graphs. Key: *, *p* < 0.05; **, *p* < 0.01 vs. control.

### Cilengitide impairs αvβ3 signaling of DAOY cells and recapitulates the β3-depletion phenotype

To mimic β3-depletion, the pharmacological effects of cilengitide, an RGD-derived compound, were examined in MB cell lines in vitro (Figure 3). The IC50 of cilengitide was determined using MTT or adhesion assays. In both assays, the effect of cilengitide was restricted to β3-expressing MB cell lines with IC50s of <2 μM and <10 μM in DAOY-WT and HD-MB03-β3+, respectively (Figure 3A). Consistent with the genetic approach, cilengitide impaired β3-downstream signaling in DAOY cells in a dose-dependent manner (Figure 3B-C). This effect resulted in anti-proliferative, anti-migratory, and anti-invasive effects at 0.5 and 2.5 μM (Figure 3D-F). Moreover, 48 h treatment with cilengitide induced cell apoptosis in DAOY-WT cells (Figure GH).

**Figure 3.**
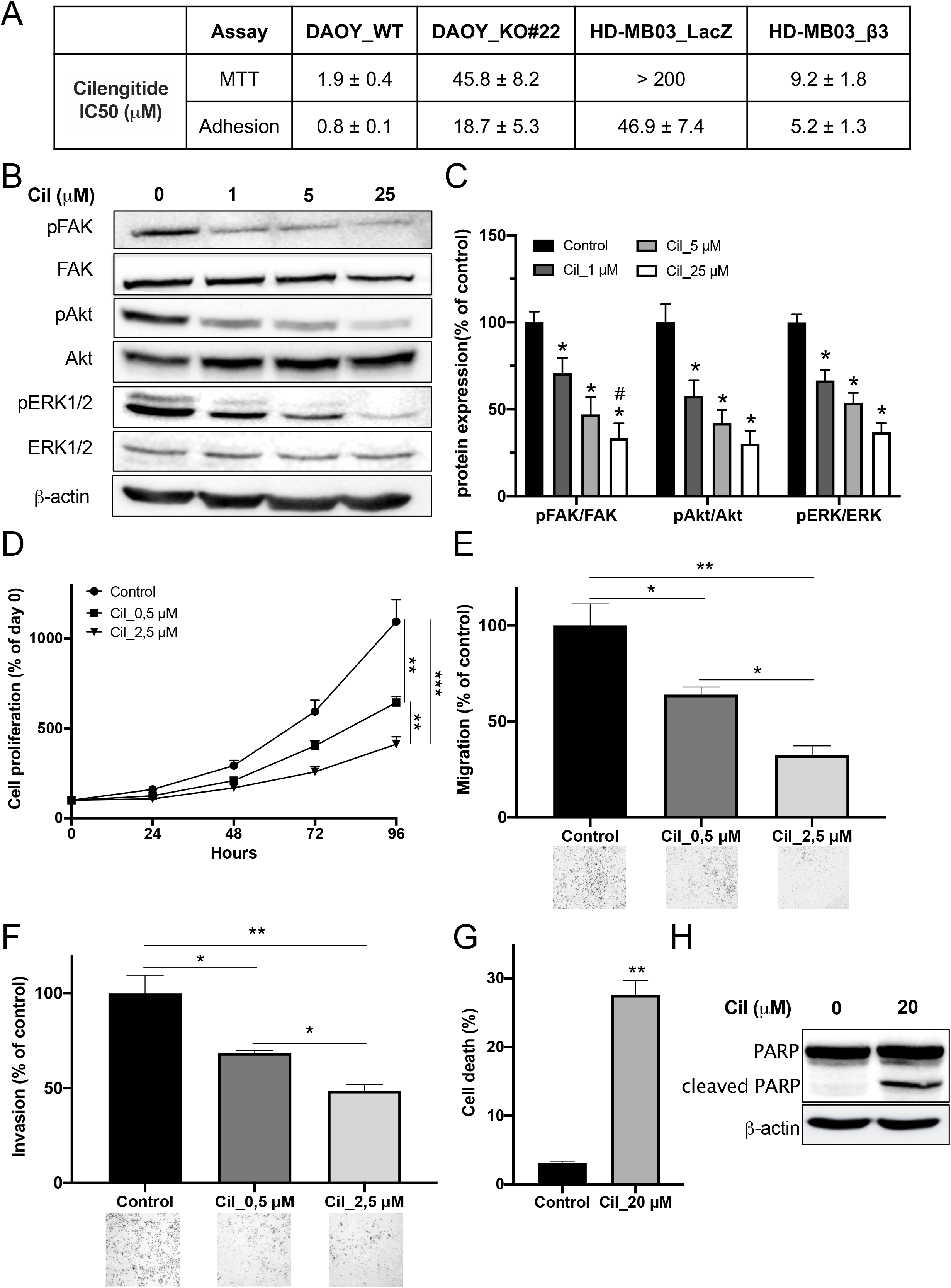
Cilengitide disrupts integrin-αvβ3 downstream signaling, decreasing cell proliferation, migration, and invasion of DAOY. (**A**) The IC50s for cilengitide determined by MTT and adhesion assays. In the MTT assay, IC50s (μM) were determined after 48 h of exposure to cilengitide. In adhesion assays, 96-well plates were coated with fibronectin (1μg/well), and cells were allowed to adhere in the presence of cilengitide for 2 h. The IC50s were determined after staining the cells with Crystal Violet (1%), resuspending them in DMSO, and absorbance measurement at 590 nm. (**B**) Representative western blots show FAK, Akt, and ERK1/2 activation in DAOY cells after 1 h of cilengitide treatment. DAOY cells were treated with 1, 5, or 25 μM cilengitide for 60 min at 37°C. Three independent experiments were performed. (**C**) Protein expression in western blots was quantified by densitometry, and the results are expressed as the percentage of untreated control cells. (**D**) The proliferation of DAOY cells treated with cilengitide (0.5 or 2.5 μM) for 96 h. The proliferation rates are expressed as the percentage of day 0. Three independent experiments were performed, and data are represented as mean ± SEM. Key: **, *p* < 0.01; ***, *p* < 0.001. (**E**) The migration of DAOY cells treated with cilengitide for 12 h. Serum-starved cells were treated with cilengitide (0.5 or 2.5 μM) and allowed to migrate for 12 h. Migrations were performed using Boyden chamber assays, and the results are expressed as the percentage of control cells. Three independent experiments were performed, and data are presented as mean ± SEM. Key: *, *p* < 0.05; **, *p* < 0.01. (**F**) An invasion assay of DAOY cells treated with cilengitide for 12 h. Serum-starved cells were treated with cilengitide (0.5 or 2.5 μM) and allowed to invade for 12 h. Invasions were conducted using Matrigel-coated Boyden chamber assays, and results are expressed as the percentage of control cells. Three independent experiments were performed, and data are presented as mean ± SEM. Key: *, *p* < 0.05; **, *p* < 0.01. (**G**) Cell death was determined by the PI staining method. Cells were treated with cilengitide (20 μM) for 48 h, then non-adherent and adherent cells were collected, and cell viability was assessed. Three independent experiments were performed, and data are presented as mean ± SEM. Key: **, *p* < 0.01 vs. control. (**H**) The effect of cilengitide on PARP cleaveage. DAOY cells were treated with cilengitide (20 μM) for 48 h. Cell lysates were analyzed by western blot with an anti-PARP antibody. Three independent experiments were performed, with representative blots shown.

### In vivo anti-tumor effects of cilengitide are restricted to αvβ3-expressing MB

Cilengitide’s efficacy was next evaluated in orthotopic models of DAOY-WT, DAOY-KO, and HD-MB03 tumors to investigate its relevance for integrin-αvβ3-expressing MBs. Cilengitide administration delayed tumor growth only in mice bearing DAOY-WT tumors (Figure 4A-B). This anti-tumor effect resulted in a 181% increase in mouse survival in the DAOY-WT model (Figure 4C, left panel). Nonsignificant effects were observed in DAOY-KO and HD-MB03 tumors, demonstrating cilengitide’s specificity for tumoral integrin-αvβ3 (Figure 4A-C). Due to its anti-angiogenic function, cilengitide administration resulted in a 60% decrease in intra-tumoral blood vessels in each model (Figure 4E). However, a decrease in Ki67 nuclei was observed only in DAOY-WT tumors, consistent with cilengitide’s anti-tumor effect (Figure 4D, left panel).

**Figure 4.**
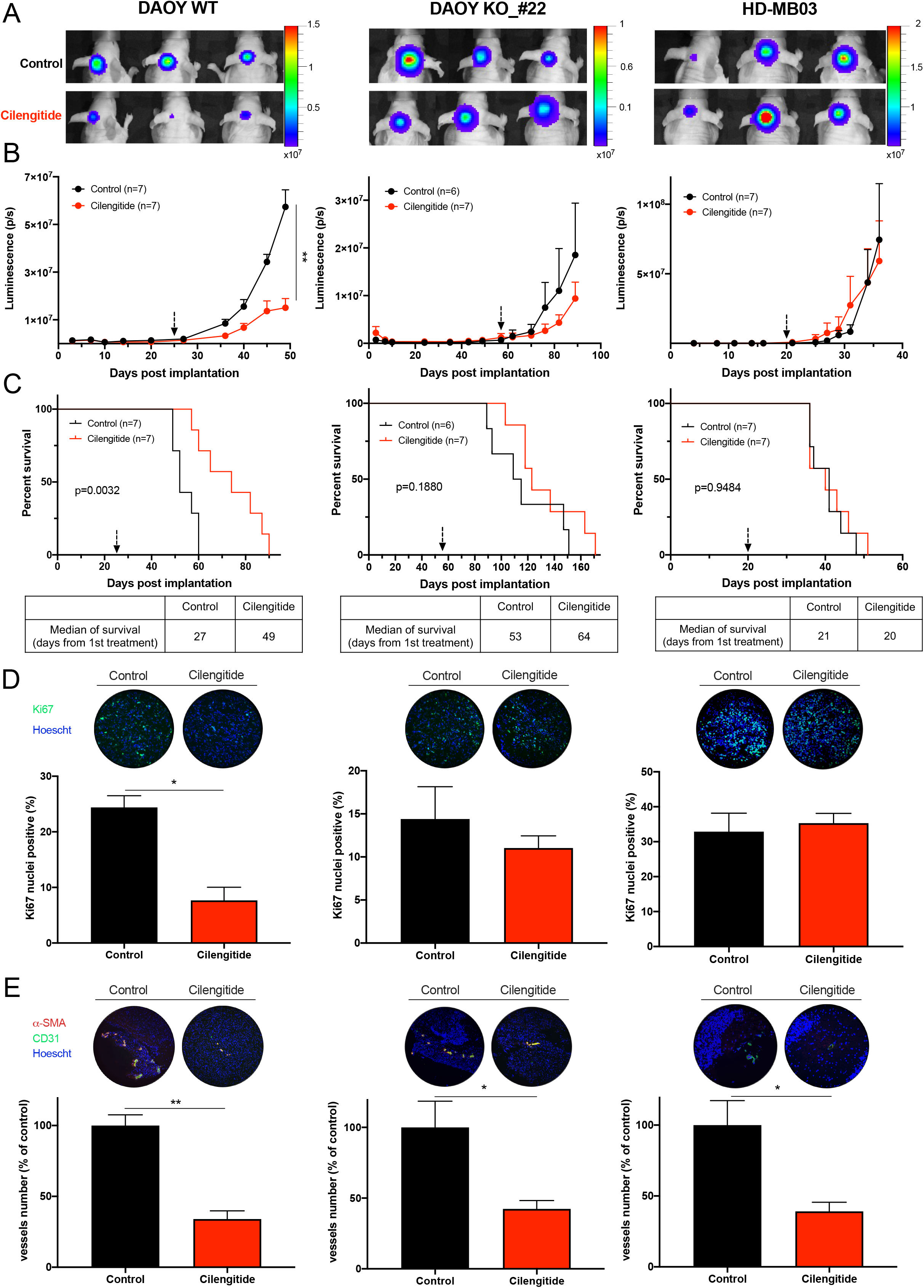
Anti-tumor effects of Cilengitide are restricted to integrin-αvβ3-positive tumors. (**A**) Representative images of the BLI signal intensity 49 days (DAOY_control, left), 89 days (DAOY-KO#22, middle), and 36 days (HD-MB03_WT, right) after implantation, corresponding to the time of first death. Three mice are shown per condition. (**B**) Tumor growth of DAOY_control (left panel), DAOY-KO#22 (middle panel), and HD-MB03_WT (right panel) as measured by luciferase activity. Photon flux was quantified and analyzed using the IVIS imaging system. Key: **, *p* < 0.01 vs. control. (**C**) Survival curves of mice orthotopically implanted with DAOY_control (left), DAOY KO_#22 (middle), and HD-MB03 (right) cells in the cerebellum and treated with cilengitide. Day 0 corresponds to tumor implantation, and the black arrow indicates the start of treatment. Mice were treated with 300 μg cilengitide three times a week. The median survival (from the start of treatment) is shown at the bottom of the graphs. The *p*-value is indicated in the graph (Log-rank test). Key: **, *p* < 0.01 vs. control. (**D**) Ki67 labeling of DAOY_control (left panel), DAOY-KO_#22 (center panel), and HD-MB03_WT (right panel) tumor sections. Representative images of Ki67 immunolabeling (green) and Hoechst33342 nuclear DNA counterstaining (blue) are shown at the top of the graphs. Key: *, *p* < 0.05 vs. control. (**E**) Vasculature labeling of DAOY_control (left panel), DAOY-KO_#22 (middle panel), and HD-MB03_WT (right panel) tumor sections. Vessels were identified by CD31 and αSMA immunofluorescent staining (green and magenta, respectively). The number of vessels (CD31+ and αSMA+) was measured, and results are expressed as the percentage of control conditions. Key: *, *p* < 0.05; **,*p* < 0.01 vs. control.

### SPECT-MRI imaging as a dual-modality strategy for measuring integrin-αvβ3 expression

Cilengitide’s restricted efficacy for αvβ3-positive tumors highlights the need for patient stratification. As a key component of personalized medicine, nuclear imaging is a relevant strategy to answer this issue. Therefore, we investigated the ability of ^99m^Tc-RAFT-RGD, a radioligand targeting integrin-αvβ3, to noninvasively evaluate its expression in DAOY and HD-MB03 orthotopic tumors (Figure 5). SPECT-MRI acquisitions showed that ^99m^Tc-RAFT-RGD uptake was readily observable in DAOY tumors, while no signal was found in HD-MB03 tumors (Figure 5A). Tumor-to-cerebellum ratios were significantly higher in DAOY tumors than in HD-MB03 tumors (*p* < 0.010; Figure 5B). Autoradiography of tumor slices was consistent with SPECT imaging quantification, with higher tumor-to-cerebellum ratios in DAOY tumors (Figure 5C-D). These results suggest that radiotracers targeting αvβ3, such as ^99m^Tc-RAFT-RGD, could be used as companion markers to identify responders before cilengitide administration.

**Figure 5.**
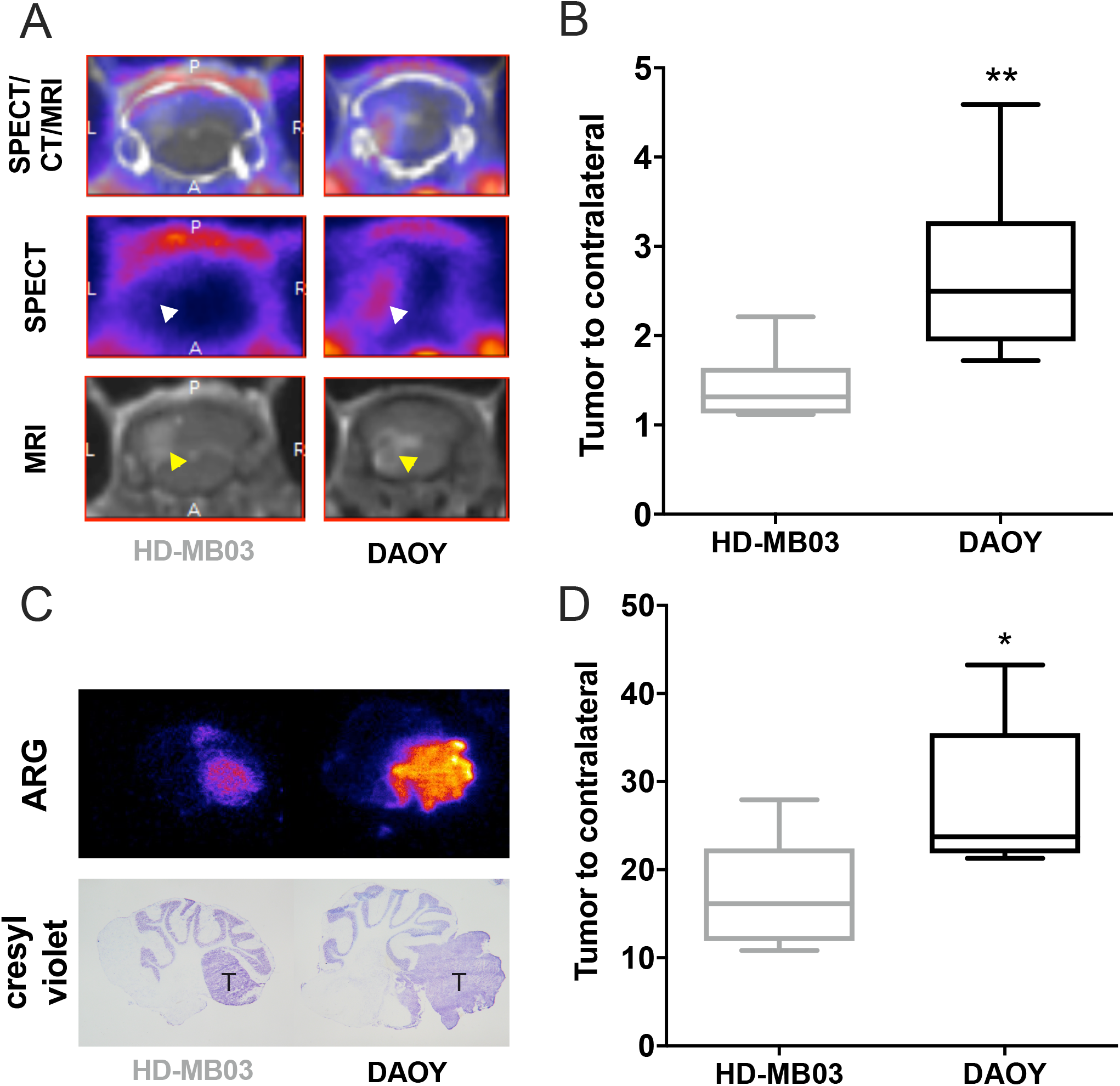
In vivo measurement of integrin-αvβ3 expression using SPECT/MRI imaging dual-modality. (**A**) Representative coronal views of MRI (bottom), SPECT (center), and fused SPECT/CT/MRI images of HD-MB03 (left) and DAOY (right) tumors at 1 h post-injection of ^99m^Tc-RAFT-RGD. (**B**) In vivo quantification of ^99m^Tc-RAFT-RGD tumor uptake from SPECT images. Results are expressed as tumor-to-cerebellum ratios. Key: **, *p* < 0.01 vs. HD-MB03. (**C**) Autoradiography on 20 μm tumor-containing cerebellum slices (top) and staining of adjacent slices by Crystal Violet. (**D**) Autoradiographic image quantification of ^99m^Tc-RAFT-RGD uptake in HD-MB03 and DAOY tumor lesions. Results are expressed as tumor-to-cerebellum ratios. Key: *, *p* < 0.05 vs. HD-MB03.

### Integrin-αvβ3 is a relevant target for radioresistant MB

Because radioresistance is an important issue in brain tumor therapy, αvβ3-integrin expression was examined in radioresistant DAOY and HD-MB03 cells generated after multiple X-ray irradiations (Figure 6). Cell viability was 60% higher for radioresistant DAOY and HD-MB03 (DAOY -RR/HD-MB03-RR) than for naïve cells after a single irradiation (Figure S5). The mRNA analysis performed in these cells showed a significant increase in *ITGB3* expression in the radioresistant cells (Figure 6A). This observation was confirmed at the protein level with higher expression in radioresistant cells (Figure 6B).

**Figure 6.**
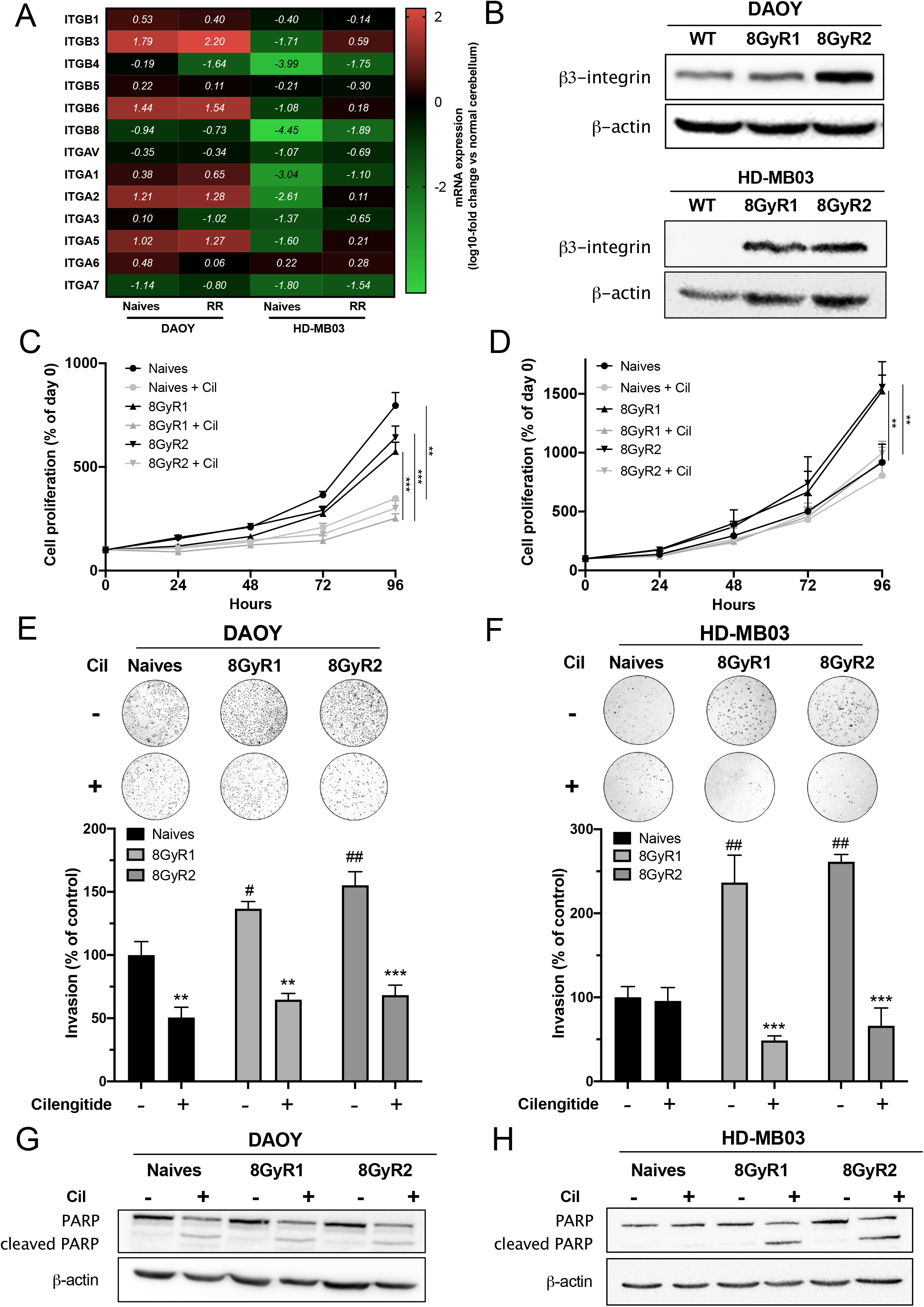
Radioresistant MBs overexpress integrin-αvβ3 and are sensitive to cilengitide. (**A**) A heatmap showing integrin gene expression in naïve vs. radioresistant DAOY and HD-MB03 cells. Integrin expression was quantified by RT-qPCR. Results are expressed compared to normal cerebellum (log_10_-fold change). The means of three independent experiments are presented. (**B**) Representative western blots of β3-integrin protein expression in naïve vs. radioresistant DAOY (top) and HD-MB03 (bottom) cells. Three independent experiments were performed. (**C**-**D**) The proliferation of naïve vs. radioresistant DAOY (C) and HD-MB03 (D) cells treated with cilengitide (2.5 μM) for 96 h. Proliferation rates are expressed as the percentage of day 0. Three independent experiments were performed, and data are presented as mean ± SEM. Key: **, *p* < 0.01; ***, *p* < 0.001. (**E-F**) Invasion assays of naïve vs. radioresistant DAOY (E) and HD-MB03 (F) cells treated with cilengitide for 12 h and 24 h, respectively. Serum-starved cells were treated with cilengitide (2.5 μM) and allowed to invade for 12 h (DAOY) or 24 h (HD-MB03). Invasions were conducted using Boyden chambers coated with Matrigel, and the results are expressed as the percentage of control naïve cells. Three independent experiments were performed, and data are presented as mean ± SEM. Key: **, *p* < 0.01; ***, *p* < 0.001 vs. untreated cells; #, *p* < 0.05; ##, *p* < 0.01 vs. naïve cells. (**G-H**) Effect of cilengitide on PARP cleavage. Naïve or radioresistant DAOY (G) and HD-MB03 (H) cells were treated with cilengitide (20 μM) for 48 h. Cell lysates were analyzed by western blot with an anti-PARP antibody. Three independent experiments were performed, with representative blots shown.

The effect of cilengitide was further investigated in vivo in both models. It decreased cell proliferation of both naïve and radioresistant DAOY cells. However, its effect was limited to radioresistant HD-MB03 cells expressing integrin-αvβ3 (Figure 6C-D). Invasion assays performed with these cells showed a significantly higher invasion potential for radioresistant cells than for naïve cells (Figure 6E-F). Cilengitide decreased the number of invasive cells in the DAOY WT and radioresistant populations but only in the HD-MB03 radioresistant population. Cilengitide induced cell apoptosis in these cells, as shown by PARP cleavage after 24 h of treatment (Figure 6G-H). Altogether, these results suggest that αvβ3-integrin is a potential therapeutic target for radioresistant MB.

## DISCUSSION

MB is the most frequently diagnosed primary brain tumor in children (16,17). Its standard treatment protocol comprises maximal safe resection followed by craniospinal radiation and adjuvant therapy. However, approximately 30% of children with MB experience disease relapse, which is almost always fatal (18). Therefore, recurrent MB is a major therapeutic challenge. Given the apparent need to improve therapeutic strategies, considerable efforts have been made to understand the biology of brain tumors. The low survival rates of brain tumors, including MB and GBM, are at least partly due to extensive brain tissue invasion.

Invasion is finely regulated and is mainly controlled by cancer and stromal cell interactions (19,20). In addition to invasive features, GBM and MB show marked tumor cell proliferation and increased angiogenesis. All these processes have mainly been studied in GBM, where integrins have been shown to be widely expressed and to play fundamental roles (21–23). Therefore, integrins have become attractive target candidates for therapeutic intervention in GBM. Among them, the pro-angiogenic αvβ3 was the first to be found to be abundantly expressed in high-grade brain tumors (23).

Integrin-αvβ3 belongs to the integrin subtypes that recognize the tripeptide RGD sequence found in many ECM proteins, including fibronectin or vitronectin. Many studies have highlighted the role of αvβ3 in sustaining GBM’s high proliferative, migrative, and invasive properties and promoting angiogenesis (24–26). Nevertheless, the expression and the role of integrin-αvβ3 have been poorly investigated in MB. In 2005, Lim et al. reported moderated integrin-αvβ3 expression in five MB patients (27). In our study, we detected significant integrin-αvβ3 expression in 20% of patient-derived samples and showed its importance for a subpopulation of MB patients. The restricted integrin-αvβ3 expression found in MB-derived samples was confirmed in MB cell lines, of which only two of five showed *ITGB3* overexpression.

DAOY and ONS-76, belonging to SHH subgroups, expressed integrins in contrast to D458 and HD-MB03 (group 3) and CHLA (group 4). These results suggest that integrin-αvβ3 may be restricted to a molecular subset of MBs. Studies should be performed to confirm this observation. Next, we investigated the role of integrin-αvβ3 in MB by genetically depleting its β3 subunit in DAOY cells and overexpressing its β3-subunit by lentiviral transduction in DAOY-KO and HD-MB03 cells. While integrin-αvβ3 has long been considered expressed in neo-vessels, IHC analysis of GBM samples showed an important expression by tumor cells rather than angiogenic cells, promoting GBM cell proliferation, migration, and invasion (23). In both cell lines, integrin-αvβ3 was a major player in these three processes. As orthotopic experiments showed, depleting the β3-subunit in two independent DAOY clones strongly reduced their tumorigenic potential. To a lesser extent, similar results were observed in HD-MB03 cells. These results support the original role of integrin-αvβ3 in MB tumorigenesis.

Since its discovery, the core integrin-binding domain RGD in fibronectin has attracted much attention in the field of anti-cancer therapies. Among the various RGD-containing peptides developed to impair tumorigenesis, cilengitide was identified as a selective αvβ3 and αvβ5 inhibitor (28–30). It showed anti-angiogenic, cytotoxic, and anti-invasive activities in preclinical GBM models. Subsequent phase III CENTRIC and phase II CORE clinical trials showed no significant effects on OS in newly diagnosed GBM (12,13). Nevertheless, in the CORE study, high αvβ3 levels in tumor cells were associated with improved OS. This finding suggests that cilengitide’s efficiency was mediated by its action on αvβ3-positive tumor cells rather than angiogenic cells.

Here, we evaluated cilengitide in our MB models. The MTT assay showed that cilengitide did not affect HD-MB03 cells, while it strongly decreased the proliferation and viability of DAOY and HD-MB03-β3 overexpressing cells. The DAOY-KO cells were found to be only slightly affected by cilengitide treatment, with an IC50 >20-fold higher than that of DAOY-WT cells. This difference could be due to αvβ5 expression in DAOY-WT and -KO cells. Cilengitide impaired downstream integrin-αvβ3 signaling in a dose-dependent manner, decreasing the phosphorylation of FAK, Akt, and ERK1/2. These molecular effects led to significant anti-proliferative, anti-migratory, and anti-invasive effects in DAOY cells.

Cilengitide was subsequently studied in vivo in orthotopic DAOY-WT, DAOY-KO, and HD-MB03 models. As expected, treatment with cilengitide significantly reduced blood vessel density in the three models, as cilengitide has been generally described as a highly potent inhibitor of angiogenesis. Cilengitide’s anti-tumor effect was limited to high integrin-αvβ3 expressing tumors (DAOY), with an 181% increase in median survival. This finding highlights the importance of stratifying patients prior to cilengitide administration and supports data from the CORE study conducted on GBM patients. Non-invasive molecular imaging could be a valuable tool to confirm the existence of the target and track its expression to monitor tumor response.

In recent decades, several αvβ3-targeting radiotracers have been developed and investigated for clinical translation (10). Most are based on the tripeptide RGD due to its high affinity and specificity for integrin-αvβ3. Preclinical and clinical studies have found no physiological brain uptake of different RGD peptides. However, clinical studies showed significant uptake in GBM patients (10). In our study, we used a ^99m^Tc-radiolabeled tetrameric RGD-based peptide (RAFT-RGD) previously validated for imaging and internal vectorized therapy in preclinical GBM models (31). Consistent with our results with cilengitide, ^99m^Tc-RAFT-RGD uptake was higher in DAOY than in HD-MB03 tumors.

The combination of SPECT and MRI in these studies paves the way for nuclear imaging use in MB patients. Our results with radioresistant MB suggest that integrin-αvβ3 is expressed in radioresistant tumors (32). Therefore, SPECT/MRI imaging in recurrent MB could be considered to determine eligibility for cilengitide treatment. Indeed, accumulating evidence supports the role of integrin-αvβ3 in resistance to conventional therapies. Other studies have reported integrin-αvβ3 expression in radioresistant gliomas and prostate cancer tumors (32,33). If integrin-αvβ3 expression is an important determinant of cilengitide efficacy, several factors, such as redundancy or compensatory mechanisms, could explain its limited efficacy in the clinic.

The remarkable advances in nuclear medicine, especially targeted therapies, may provide new tools for treating recurrent MB. Most therapeutic radiopharmaceuticals are labeled with β-emitting isotopes. These particles penetrate only a few millimeters into tissue and allow irradiation of cells within a limited radius, resulting in tumor cell death while sparing surrounding healthy tissue. Commonly used β-emitters include lutetium-177 (^177^Lu-), which has a half-life of 6.7 days and relatively high β-emission (0.497 MeV). These characteristics offer the advantage of delivering a high dose and achieving prolonged tumor irradiation. In addition, a unique feature of radionuclides is that they can exert a “cross-fire” effect that destroys neighboring cells. Several ^177^Lu-radiolabeled RGDs have been studied in preclinical GBM models. One, the ^177^Lu-NOTA-EB-RGD, was found to completely eradicate tumor growth from αvβ3-expressing patient xenografts (34). Although all these compounds require further clinical investigation, they may have great potential for integration into combined therapies for brain tumors, including MBs. Future perspectives of our study include the evaluation of ^177^Lu-RAFT-RGD in mouse MB models, which could be the beginning of the era of theranostics in these tumors.

## Supplementary data

**Table S1.**
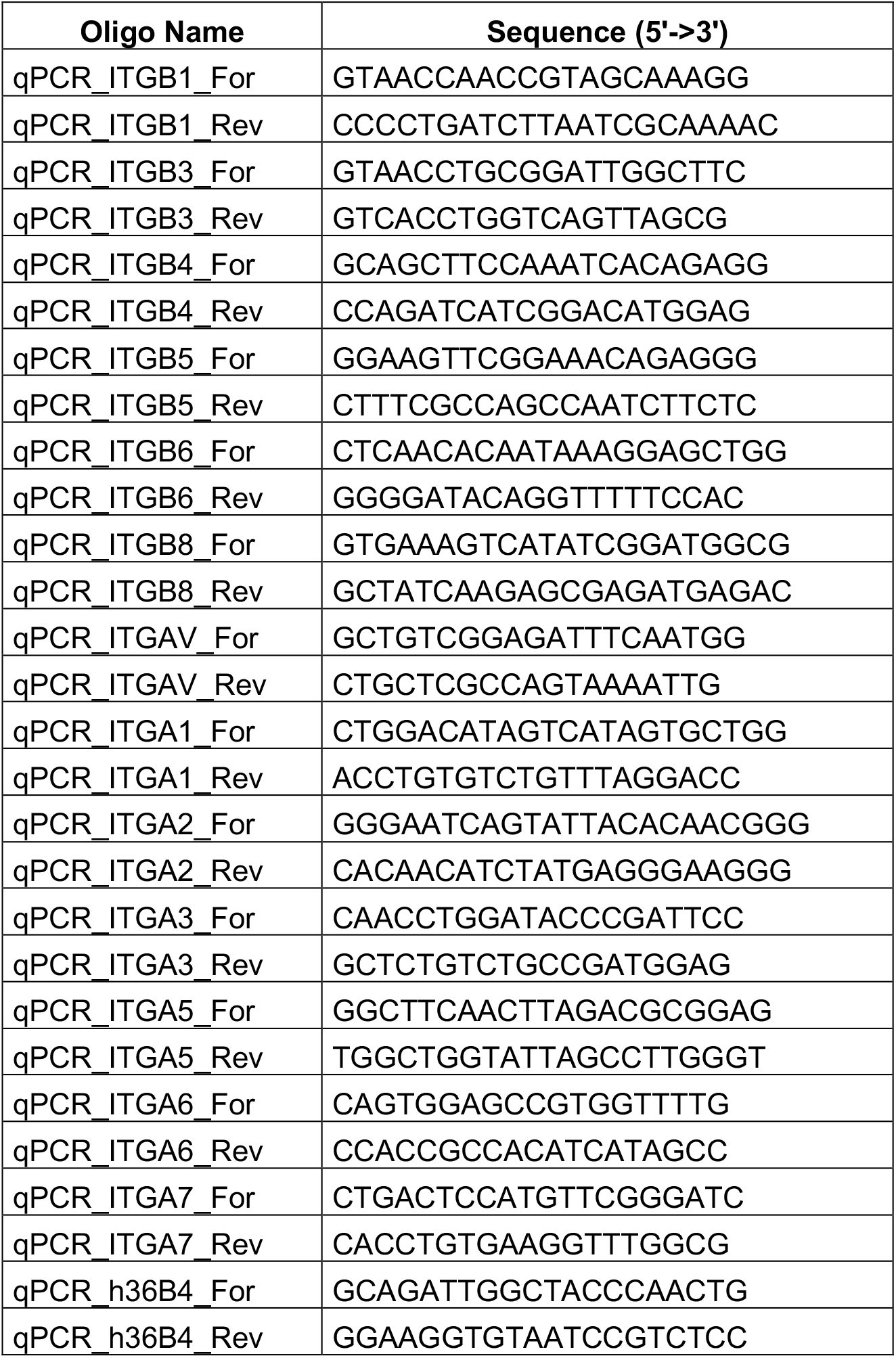
List of the primers used for qPCR analysis in this study.

**Figure S2.**
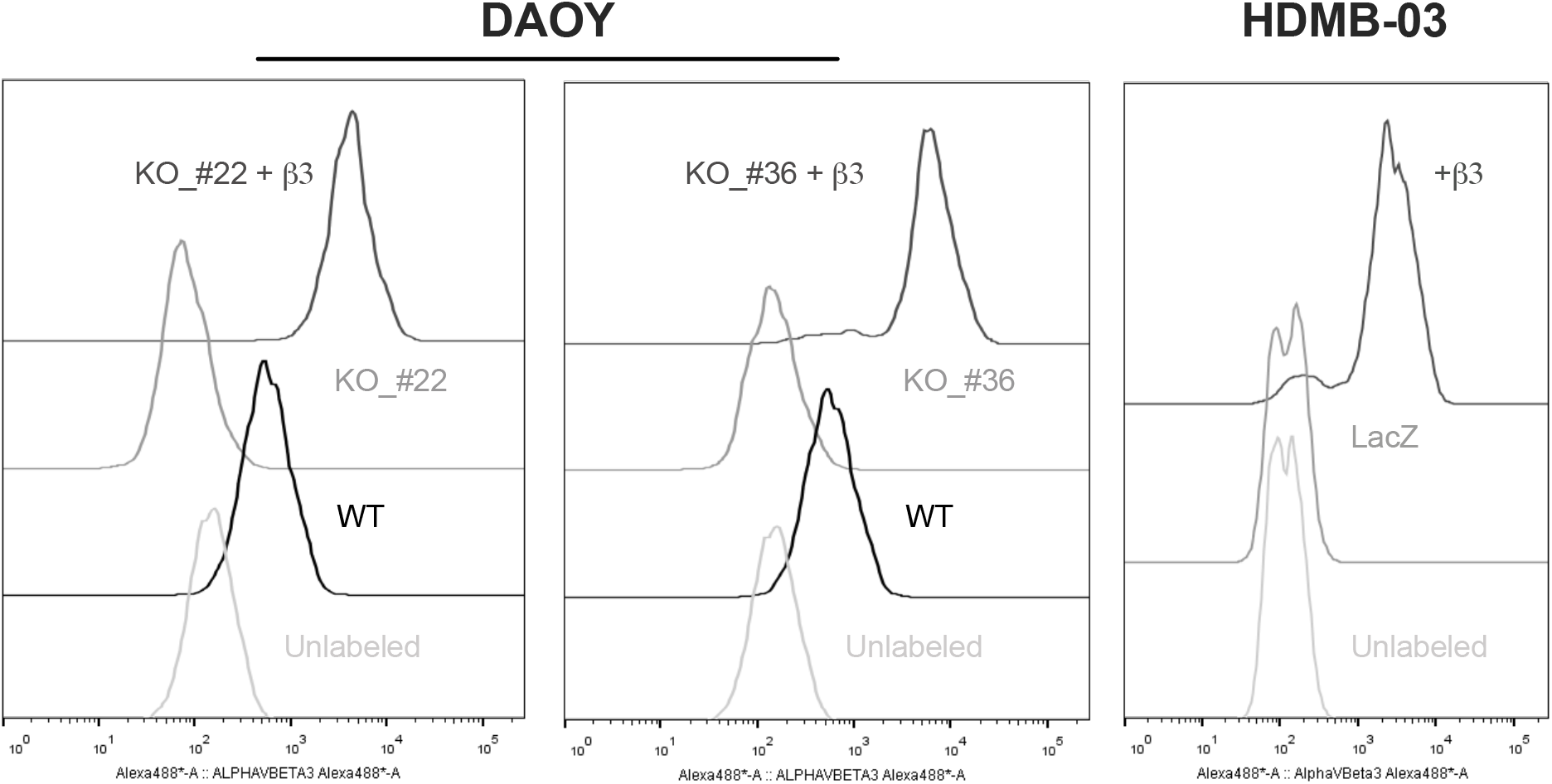
Measurement of integrin-αvβ3 expression in MDB-derived cell lines performed by FACS. DAOY- and HD-MB03-derived cells were seeded in 6-well dishes for 24h. After detachment with accutase, cells were collected and incubated with an anti-αvβ3 antibody (ab190147, Abcam®) followed by a goat antimouse secondary antibody coupled to AlexaFluor™488. Data were analyzed per sample using a BD FACSMelody™ cytometer. Data were analyzed using FlowJo™ software.

**Figure S3.**
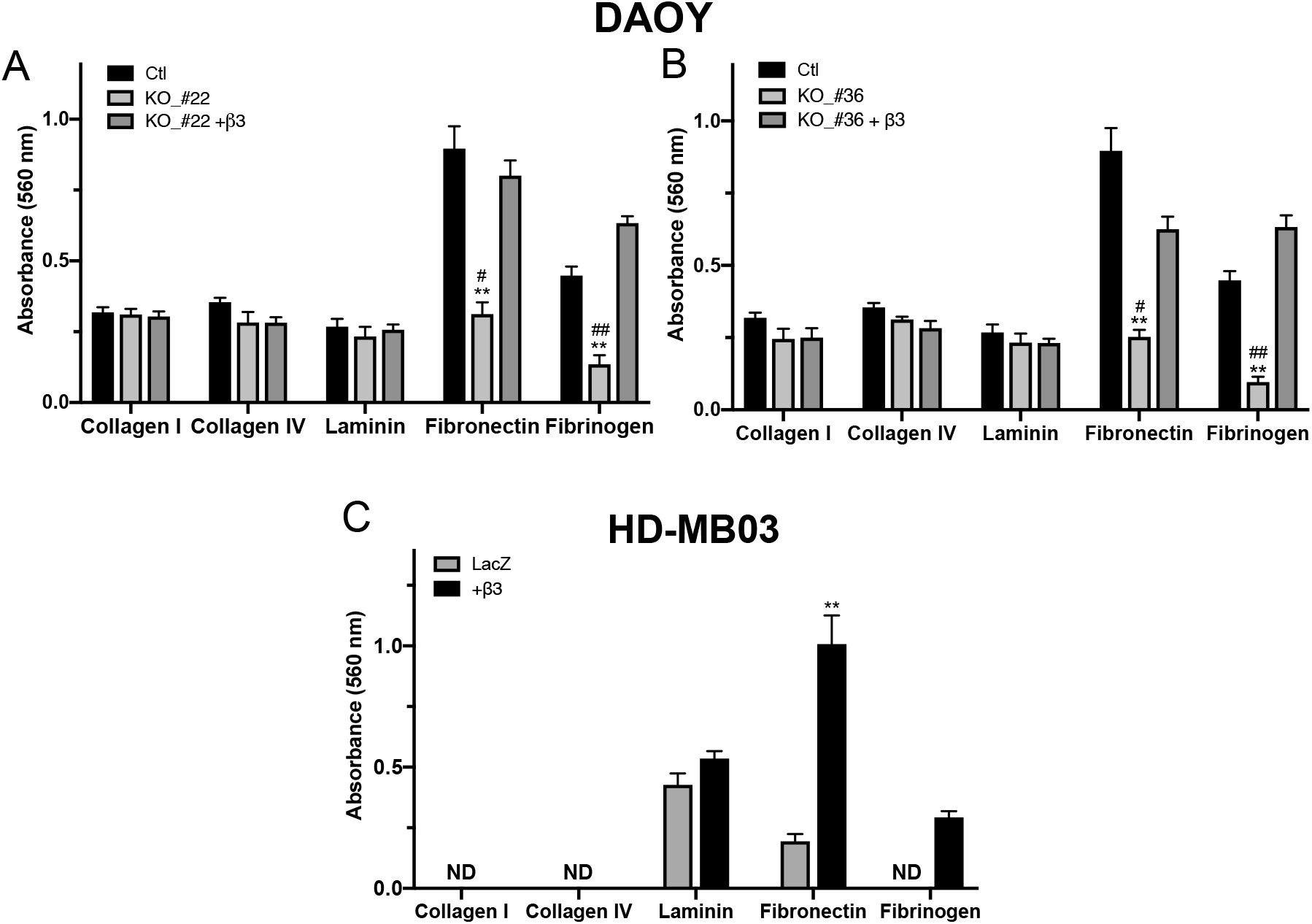
ECM-mediated cell adhesion assay using crystal violet staining. DAOY-derived (**A**-**B**) and HD-MB03-derived (**C**) cells were allowed to attach to different ECM proteins coated on 48-wells plates for 30 min. Adherent cells were stained with crystal violet, solubilized with DMSO and quantified at OD560 nm. ** p<0.01 vs DAOY_Ctl or HD-MB03_LacZ; # p<0.05, ## p<0.01 vs DAOY_β3-overexpressing cells.

**Figure S4.**
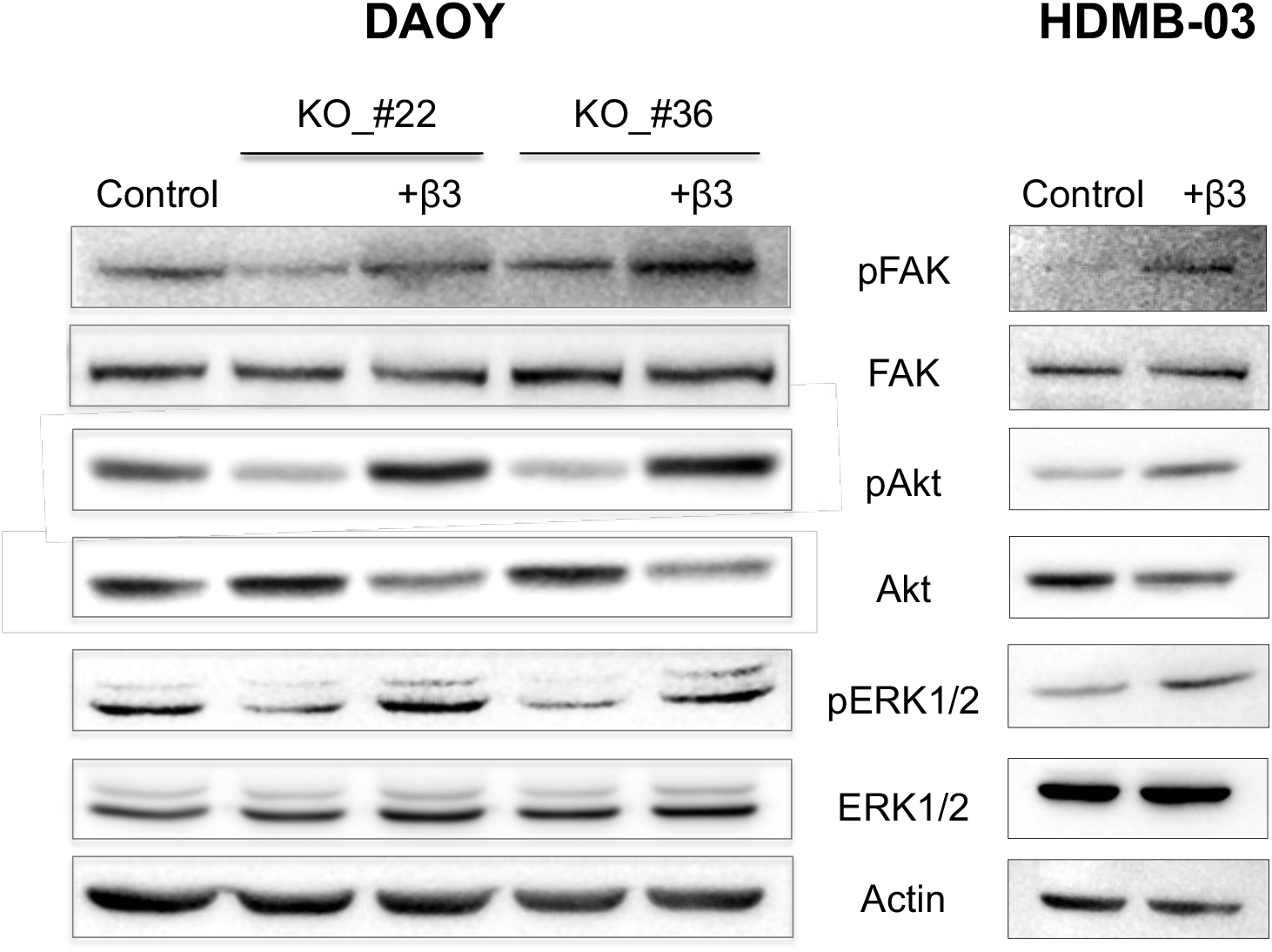
Western-blot analysis of integrin-αvβ3-downstream pathways. Relative protein contents of pFAK/FAK, pAkt/Akt and pERK1/2/ERK were determined in DAOY-derived and HD-MB03-derived cells. Actin acted as a protein-loading control and blots are representative of three independent experiments.

**Figure S5.**
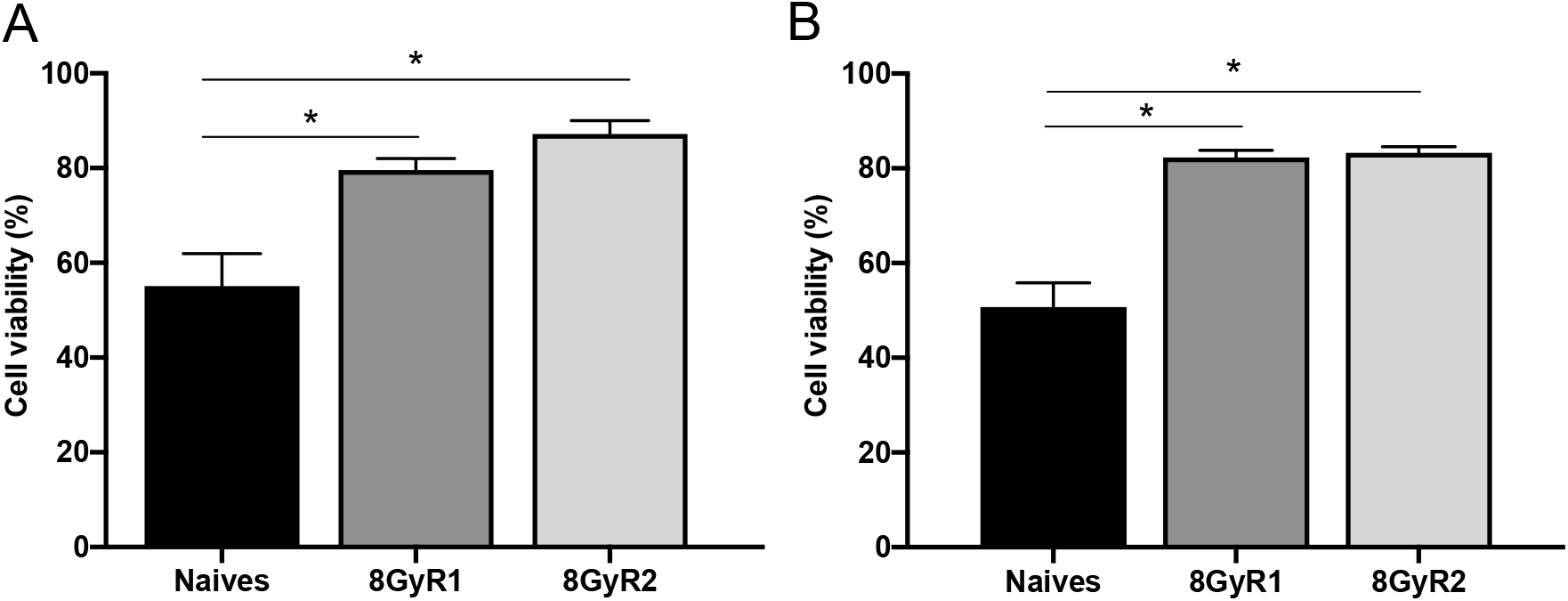
Viability of MB-derived cells after 10 cycles of X-Ray irradiation (8 Gy). Two populations of naïve DAOY (A) or HD-MB03 (B) cells have been irradiated weekly, for 10 weeks. An eleventh irradiation was conducted to perform viability tests. Viability assay was assessed by PI measurement (FACS analysis). * p<0.05 vs indicated group.

